# PTSD Biomarker Database: Deep Dive Meta-database for PTSD Biomarkers, Visualizations, and Analysis Tools

**DOI:** 10.1101/547901

**Authors:** Daniel Domingo-Fernández, Allison Provost, Alpha Tom Kodamullil, Josep Marín-Llaó, Heather Lasseter, Kristophe Diaz, Nikolaos P. Daskalakis, Lee Lancashire, Martin Hofmann-Apitius, Magali Haas

**Affiliations:** Department of Bioinformatics, Fraunhofer Institute for Algorithms and Scientific Computing, Sankt Augustin 53754, Germany; Cohen Veterans Bioscience, 1 Broadway, Cambridge, MA 02142, United States

**Keywords:** Biomarkers, Post-traumatic stress disorder, Metadatabase, Database, Bioinformatics

## Abstract

The PTSD Biomarker Database (PTSDDB) is a database that provides a landscape view of physiological markers being studied as putative biomarkers in the current post-traumatic stress disorder (PTSD) literature to enable researchers to quickly explore and compare findings. The PTSDDB currently contains over 900 biomarkers and their relevant information from 109 original articles published from 1997 to 2017. Further, the curated content stored in this database is complemented by a web application consisting of multiple interactive visualizations that enable the investigation of biomarker knowledge in PTSD (e.g., clinical study metadata, biomarker findings, experimental methods, etc.) by compiling results from biomarker studies to visualize the level of evidence for single biomarkers and across functional categories. This resource is the first attempt, to the best of our knowledge, to capture and organize biomarker and metadata in the area of PTSD for storage in a comprehensive database that may, in turn, facilitate future analysis and research in the field.

**Database URL:** https://ptsd.scai.fraunhofer.de

## Introduction

Post-traumatic stress disorder (PTSD) is a common psychiatric disorder that occurs in some individuals after a traumatic event (Pietrzak, *et al*., 2012) and is diagnosed by mental health professionals based on the presentation of four symptom clusters - intrusions, avoidance, negative cognitions/mood, and hyperarousal (American Psychiatric Association, 2013). PTSD pathophysiology is complex and affects multiple interconnected biological systems that regulate mental and physical health functions and are associated with PTSD’s clinical heterogeneity and diverse comorbidity profiles (Pitman, *et al*., 2012; Kang, *et al*., 2015; Passos, *et al*., 2015; Bina and Langevin, 2018; Daskalakis, *et al*., 2018; Speer, *et al*., 2018).

An extensive amount of research in PTSD has explored the utility of physiological markers as being discrete biomarkers of this disorder; however, no such putative biomarkers of PTSD has been identified to date. In the PTSD literature, the types of physiological markers most commonly studied include neuroimaging and psychophysiological measures, behavioral and neurocognitive read-outs, and analytes measured in peripheral biofluids, such as blood and saliva at baseline or after psychological challenge (Michopoulos *et al*., 2015; Etkin and Wager, 2007; Jovanovic and Ressler, 2010; Zoladz and Diamond, 2013; Daskalakis, *et al*., 2016). Fluid-based peripheral biomarkers may include inflammation indicators, hypothalamic pituitary adrenal axis mediators, neurosteroids, and neurotransmitters, which have functional roles both in the peripheral and central nervous system, potentially enabling biologically meaningful inference with clinical utility (Daskalakis, *et al*., 2016).

The increasing amount of biomarker studies in all disease areas is paralleled by the growing number of meta-analyses that combine data from multiple studies to systematically derive common conclusions. However, the lack of disease-specific biomarker registries impedes the harmonization and integration of results from these studies, which often remain in the form of non-structured text, figures, tables, or supplementary files. Organizing and storing this knowledge is essential to provide a comprehensive view of the biomarker landscape and to foster the discovery and development of diagnostics and treatments.

Along these lines, several biomarker databases have been recently developed that focus on specific disease domains, such colorectal cancer (Zhang *et al*., 2018), Alzheimer’s disease (Kinoshita and Clark, 2007), tuberculosis (Yerlikaya *et al*., 2017), and liver cancer (Dai *et al*., 2014). Furthermore, there are multiple resources that store and catalogue biomarker information from multiple indications, such as the Online Mendelian Inheritance in Man (OMIM) (Hamosh *et al*., 2005), the cancer biomarker database (Tamborero *et al*., 2018), and the infectious disease database (Yang *et al*., 2007). These resources illustrate how biomarker information can be curated and harmonized, and currently serve as hubs for biomarker research in their respective areas. Although biomarkers of PTSD – once identified – have the potential to critically improve patient outcomes, groups have yet to embark on similar efforts for a PTSD-specific database.

To address this, we are developing the comprehensive PTSD Biomarkers Database (PTSDDB), focusing on fluid-based biomarkers to bring together published findings within the context of study design and related results. Organizing and contextualizing published information and data is critical to understand the level of confirmatory and contradictory evidence for single biomarkers. Overall, this work is necessary to cross the translational divide between basic science discovery and clinical implementation. Considering findings for single biomarkers in tandem with details around study design may offer insights into the robustness and replicability of studies. Here, we present the first version of the PTSDDB as a biomarker database that provides a comprehensive and interactive view of results from an extensive, systematic curation effort in over 100 PTSD-focused articles.

## Materials and methods

### Curation procedure and database content

#### Corpus selection

Articles included in the PTSDDB were compiled via two routes: Recommendations and referrals from experts in the field and mining cited references from PTSD review publications. In general, publications included in this deep-dive database were original articles published from 1997 to 2017 that evaluated fluid-based biomarkers in humans, with a focus on PTSD patients versus control populations (e.g., healthy controls, trauma-exposed controls, and/or patients with psychiatric disorders or other comorbidities). Exclusion criterion included publications that did not include a PTSD population, those that included PTSD patients but in the absence of fluid-based biomarkers, or were preclinical studies.

#### Data extraction and quality assessment

The biomarker metadata information contained in the resulting 109 publications was manually extracted by five independent trained curators and added to a data model template. The spectrum of curated metadata covers many fields, including study design, demographics, study findings, assay information, and statistical methods. Ultimately, three rounds of quality control (QC) were conducted to ensure the fidelity of the metadata. In each round of QC, the metadata was reviewed by a distinct curator; if inconsistencies were found, curators worked together to reach a consensus. While there exist other QC procedures ensuring the quality of the curated data such as inter-curator agreement, these involve significant time constraints as they require two distinct curators working in parallel to extract the same information. On the other hand, the previously described QC procedure allowed us to include a larger amount of biomarker metadata while maintaining the quality of data curation. The data model template for metadata extraction was predefined based on the initial set of ten articles. However, new metadata fields to were added as necessary to accommodate new types of information being extracted from additional manuscripts.

### Database design and web application implementation

#### Database model

One challenge in data integration when curating different sources of information is organizing this knowledge according to a common schema, since this crucial point greatly influences subsequent steps of data management and analysis. In order to structure the information comprised in the 193 columns of the curation template/worksheet, we designed a data model storing this information into 17 different models (e.g., publication, biomarker, clinical study, cohort metadata etc.), which are represented as tables in the database (**Table 1**). Next, we implemented a parser of the curation template that populates the MySQL database and controls the quality of the worksheet by identifying duplicates, checking the syntax, and normalizing terms. Finally, a web application integrated the database enabling users to query, visualize, and analyze the curated content as illustrated in this manuscript.

**Table 1.** Types of information extracted from each manuscript and stored as entries in the PTSDDB. For each data category, extensive information was curated and stored as separate entries in the PTSDDB. For example, the Data Category “Biomarker” includes information on Biomarker name, HUGO ID or other acronym, gene symbol/identifier, and biomarker application (e.g., biomarkers for disease risk, patient stratification, diagnostic marker, predictive markers of disease severity or treatment response, and safety/toxicity biomarkers).

| Model/Data Category | Fields |
| --- | --- |
| Publication | PubMed identifier, authors information (e.g., names, year of publication, and geographical information) |
| Biomarker | Biomarker name, HUGO ID or other acronym, gene symbol, biomarker application |
| Time Point | Study time point (e.g., 6 weeks post-trauma, 2 years post-trauma) |
| Approach<br>Panel Details | Name, statistical method, cutoff used (e.g., *p*-value, False Discovery Rate, etc.), whether the biomarker was part of a panel, other analytes included, risk SNP, allele risk and additional notes on risk SNP |
| Numerical Summary | Statistics (i.e., mean and standard deviation [SD]) about the biomarker measurements for: primary indication, trauma exposed controls, other central nervous system and healthy controls |
| Clinical Instrument | Clinical instrument for primary indication, trauma in adult, childhood and lifetime (e.g., Clinician-Administered PTSD SCALE [CAPS], PTSD Checklist [PCL], Structured Clinical Interview for DSM-5 [SCID], etc.) |
| Clinical Study | Type of study (e.g. cross-sectional, longitudinal), timeline, challenge type, treatment response study, number of subjects per indication (e.g., trauma-exposed PTSD, trauma-exposed controls, healthy controls, other indications) |
| Indication | Name and specifics of the condition (e.g., PTSD, Childhood trauma, maternal PTSD) |
| Comorbidity | Name, comorbidity measurements (e.g., mean and SD) |
| Numerical Readouts | Statistical details of each different group included in the study (e.g., mean, SD of the biomarker for primary indication or control) |
| Inclusion/Exclusion<br>Criteria | The description of inclusion and exclusion criteria provided by each publication |
| Cohort Name and<br>Demographic Details | Details about the cohort (e.g., percent female plus the overall mean and SD in trauma-exposed PTSD vs. trauma-exposed and healthy controls, mean age and SD age of subjects) |
| Ancestry | Ancestry details for Caucasian, African American, and other ancestry of the cohort (e.g., percentage of each group included in the study) |
| Study Findings | Direction of change of the biomarker in cases versus controls as specified by the study authors (e.g., increased, no change, decreased), specific circuit changes, notes and descriptions |
| Assay | Assay details: assay brand, probe, fluid, biological substrate, assay brand, assay limit detection and measurement units |
| Assay Calculations | Mean and SD concentration (in primary indication, trauma exposed controls, healthy controls and CNS controls), sigma combined, and effect size |
| Statistical Info | Mean, SD and variance, methylation change, t and *p*-value |

#### PTSDDB implementation

The application was implemented following a model-view-controller (MVC) software architecture. The back-end is written in Python using the Django web framework technology (https://www.djangoproject.com/). Django embraces the MVC paradigm by storing the data into a MySQL relational database controlled by views that are responsible for querying the database and rendering its content to the users. The front-end renders interactive visualizations using a collection of powerful Javascript libraries: D3.js (https://d3js.org/), C3.js (http://c3js.org/), DataTables (https://datatables.net/), and DataMaps (https://datamaps.github.io/). Because the main goal of the web application is data exploration and visualization, the front-end is powered by Bootstrap, thereby retaining full compatibility with a broad range of devices (e.g., smartphones, tablets, laptops, etc.). Finally, PTSDDB is complemented with a RESTful API documented with an OpenAPI specification (https://www.openapis.org).

## Results and discussion

The PTSDDB is a comprehensive and interactive database designed to facilitate the investigation of large amounts of knowledge around PTSD physiological markers reported in the literature. The PTSDDB catalogs information on more than 900 physiological markers, enabling a broad investigation of biomarkers implicated in PTSD pathogenesis. Below, we discuss the five main pages and their applications, with the help of the most common biomarkers stored in the database (e.g., cortisol, *BDNF, FKBP5, IL-6, NR3C1*, etc.).

The first page, *“biomarkers overview”*, presents an overview of the biomarker knowledge available in the database, depicted the frequency of captured biomarkers (**Figure 1a**), the biofluids in which they were measured, the relative changes reported, and the biological substrates captured (e.g., DNA, RNA, protein). For instance, when evaluating the biological substrates used in the five most frequently captured biomarkers, we found that *BDNF* and *IL-6* are predominantly measured as a protein, *FKBP5* as mRNA, and *NR3C1* has been studied across multiple substrates; in contrast, the most common biomarker, cortisol, is measured as a blood analyte given that it - unlike the others - is a steroid hormone rather than a gene or a gene product (**Figure 2a**). Further, when examining the biological substrates in which these biomarkers are measured, cortisol and *BDNF* are most frequently assessed in serum or plasma, *FKBP5* in whole blood, *IL-6* in saliva or plasma, and *NR3C1* is equally measured in peripheral blood mononuclear cell (PBMC), blood, and saliva.

**Figure 1.**
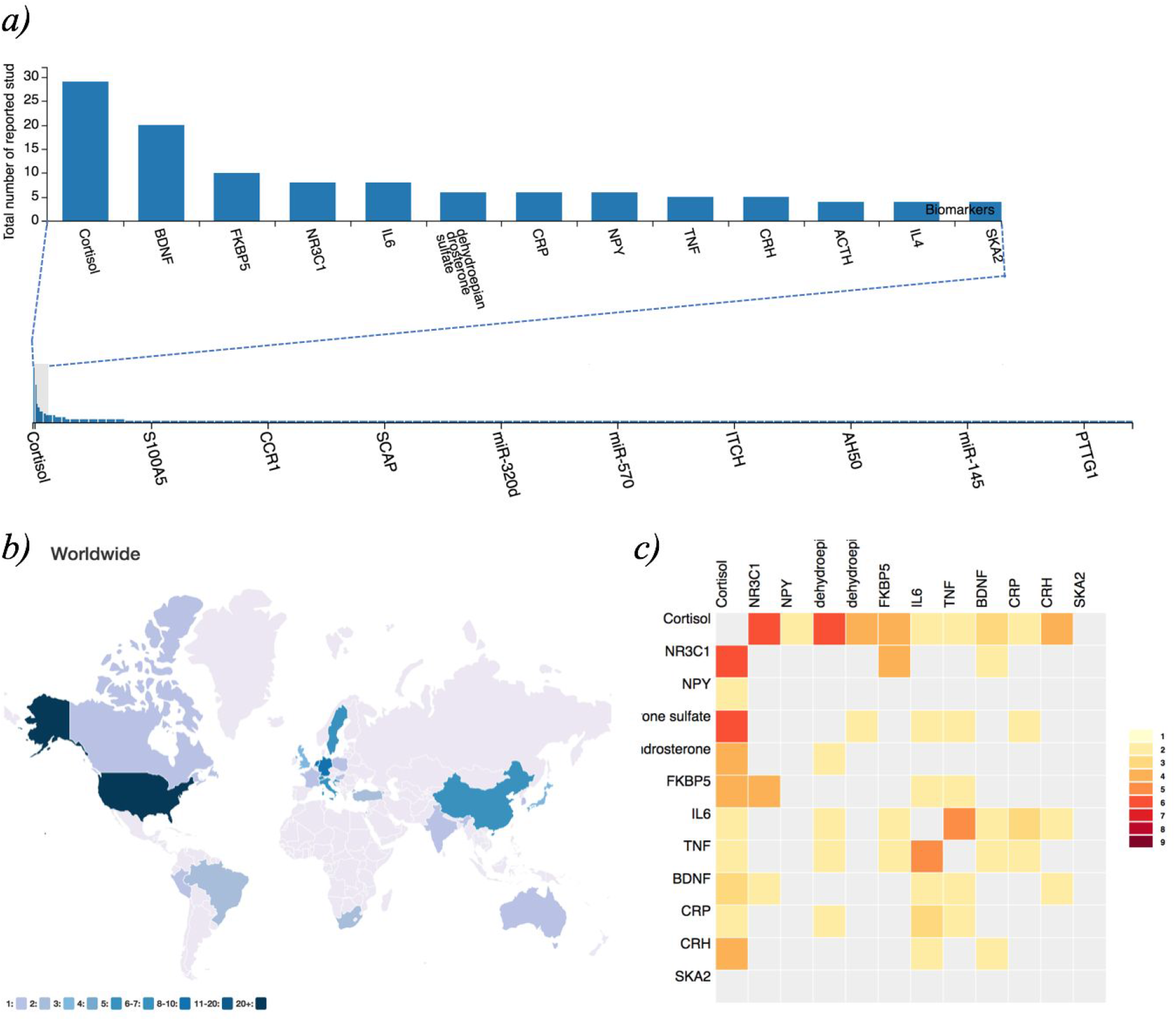
PTSDDM – Biomarker Data and Integrated Metadata: a) Frequency plot of biomarkers captured in the beta version of the PTSDDB, b) Geographical map displaying locations of institutions in the curated literature, and c) Heatmap visualization showing the frequency of individual biomarkers studied together in the same articles curated in the PTSDDB. Descriptions of these visualizations are outlined in the **Supplementary Information**, and these figures can be dynamically explored at https://ptsd.scai.fraunhofer.de/frequencies and https://ptsd.scai.fraunhofer.de/literature.

**Figure 2.**
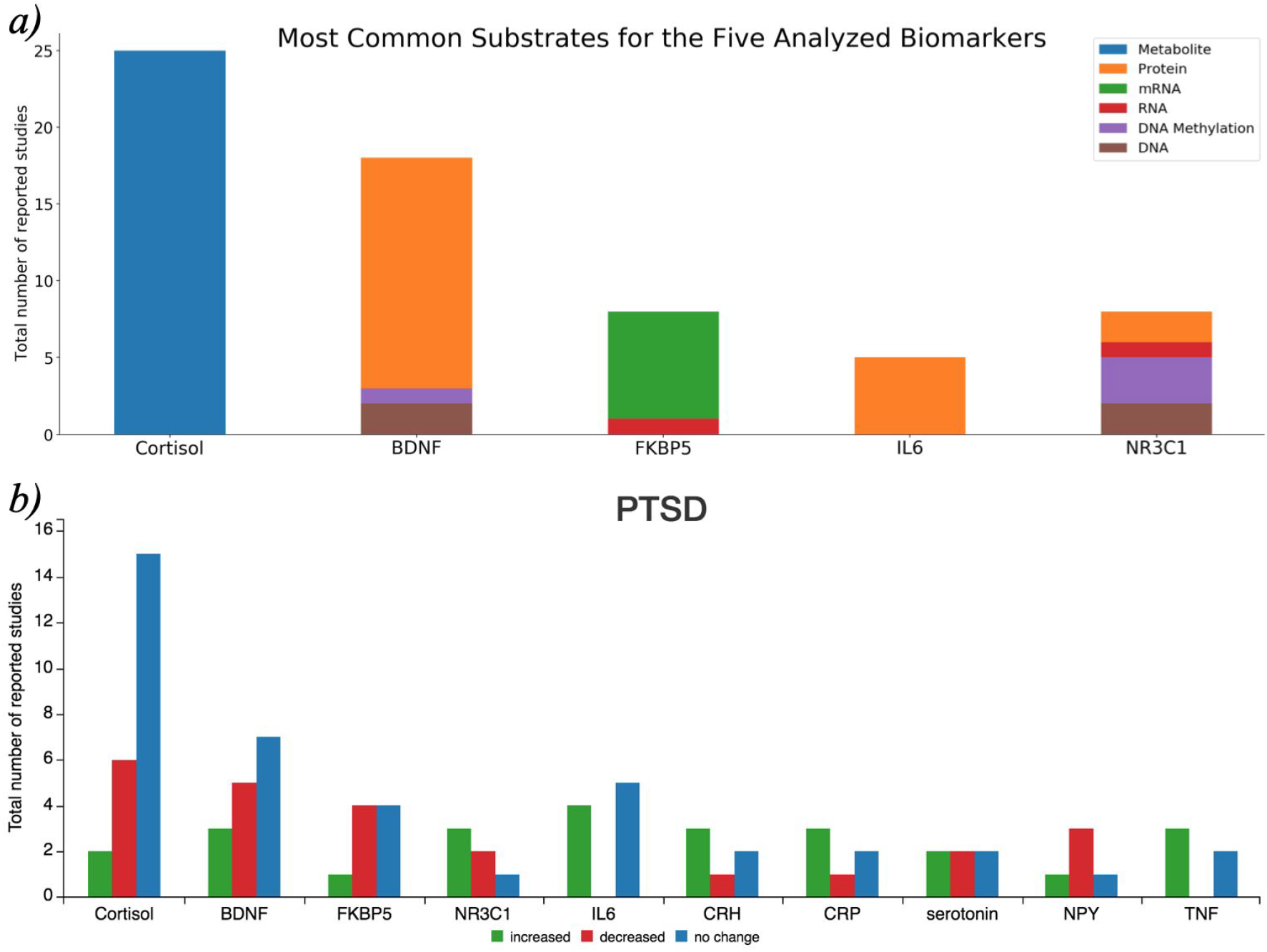
a) Biological substrates of the five most reported biomarkers in PTSDDB when they are studied as a metabolite, protein, RNA substrates. The data used to generate this figure is available through the PTSDDB API at https://ptsd.scai.fraunhofer.de/api/biological_substrate/count. b) Relative changes in the ten most common biomarkers captured in the database. This figure can be explored interactively at at https://ptsd.scai.fraunhofer.de/relative_changes.

Evaluating the accumulated evidence across studies is critical when validating biomarkers. To assist with this process, the PTSDDB provides a dedicated page, *“direction of change by biomarker”*, which contains functionalities and visualizations that summarize the directionality of biomarker findings (i.e., whether a biomarker was observed to be increased, decreased, or unchanged in cases versus controls). Moreover, the functionality of this page allows users to find articles that have reported changes in specific biomarkers of interest, evaluate the different biological substrates of origin, and compare results to determine complementary or contradictory evidence. Figure 2b summarizes the reported changes in ten of the most common biomarkers captured in this database. As a case study, we can see how there are discrepancies in findings for five of these biomarkers. Findings around *NR3C1* tend to be the most contradictory since three studies reported an increased in *NR3C1* levels, two studies reported decreased levels, and one unchanged levels in the context of in PTSD. Similarly, there exists conflicting evidence for the involvement of cortisol, *FKBP5*, and *BDNF* in PTSD, although these tend to be reported more frequently as decreased than increased (Angelucci *et al*., 2014; Bicanic *et al*., 2013; Dell’Osso *et al*., 2019; Geracioti *et al*., 2008; Gill *et al*., 2008; Inslicht *et al*., 2014; Logue *et al*., 2015; Martinotti *et al*. 2015; Stratta *et al*. 2016; Szabó *et al*. 2014; Van Zuiden *et al*., 2012; Yehuda *et al*., 2009; Yehuda *et al*., 2015; Zaba *et al*., 2015). Finally, while studies that analyzed *IL-6* seems to agree on their findings by reporting an increased concentration of this biomarker when comparing PTSD cases versus controls, there are also studies reporting no change, which was previously described in the literature (Plantinga *et al*., 2013).

Such visualizations will enable users to interact with articles and data curated in the PTSDDB, facilitating their ability to investigate and find the metadata information that might explain partially contradictory results. For instance, users may glean important information by evaluating and comparing studies around domains such as study type (cross-sectional vs longitudinal), sample size (e.g., number of subjects, controls, etc.), and experimental methods (e.g., how was the biomarker measured). Therefore, the PTSDDB includes a search functionality to contextualize the directionality of the changes based on the biological substrate where the biomarker was measured as well as a dedicated page to explore metadata information, which will subsequently be described.

While abstracts and results sections summarize the essential findings in biomarker publications, capturing the level of evidence supporting single biomarkers requires information such as sample size or statistical measurements (e.g., mean, standard deviation, etc.), and this data is often unstructured in the form of figures or supplementary files. Because the PTSDDB stores and catalogs biomarker-related metadata, one of the pages, the *“study metadata”*, enables users to access and query this information. In contrast to the previously described information, this page focuses on the sparse world of clinical metadata, which are important to compare study design. Here, users can first search for studies containing a particular biomarker and then inspect their metadata (e.g., type of study, duration, challenge, trauma, sample size, assay, etc.) for further analysis. Additionally, users can filter the studies by specific application of the putative biomarker (i.e., diagnostic, prognostic, risk, and stratification), allowing for more precise inquiries.

Biomarker research is often driven by current trends, technologies, and scientific groups that are interested in specific hypotheses related to their knowledge domain. Currently, the increasing quantity of data, information, and knowledge makes it incredibly complicated for researchers to stay abreast of all new studies published in their domain areas. Thus, it is essential to provide researchers with an overview of what biomarkers have already been investigated as well as who and where was the study conducted. This information not only allows scientists to be aware of what has been studied, but also promotes collaboration amongst those working on similar hypotheses. To provide an overview of the literature included in the database and foster new research, PTSDDB includes a page with novel visualizations, *“literature analysis”*, illustrating which biomarkers are frequently studied together; where, who, and when were the studies were conducted; or in which biological substrates the biomarkers were measured. First, a table displays the main article information: PubMed-ID, title, journal, authors, and year of publication. Second, map-based visualizations represent the geographical distribution of the analyzed articles to help identify PTSD-focused research hubs in the United States and across the globe, which may help identify collaborative opportunities for biomarker replication and validation (**Figure 1b**). Third, a histogram of the years when the articles included were published (**Supplementary Information**). Finally, two different heatmaps depict which biomarkers are frequently reported together in publications (**Figure 1c**) and which are studied in the same bio-fluid (**Supplementary Information**). By exploring this, we can investigate how the different combinations of biomarker studies were carried out, for example, the most studied biomarker pairs (e.g., cortisol and dehydroepiandrosterone sulfate cortisol and *NR3C1* were measured together in five studies) or which biological substrates were used together in clinical studies.

Since PTSDDB contains a large number of variables and metadata (**Figure 3**), it is an arduous task to implement interactive visualizations for every possible database query. Therefore, the last page, *“database content,”* contains a RESTful API (https://ptsd.scai.fraunhofer.de/swagger-ui) that exposes the database as well as a summary table of the database, providing both interactive and programmatic interfaces to query, browse, and navigate its content. The API is the gateway for researchers who are interested in data models that cannot be accessed through the interactive visualizations presented before (i.e., ancestry information, assay details and calculations, statistical information, and inclusion/exclusion criteria). This enables researchers to access specific information extracted from the study, ranging from inclusion/exclusion criteria (e.g., type of medication excluded, comorbidities excluded, etc.) to details about the equipment used in the study (e.g., machine, brand, limits of detection, etc.). Furthermore, users can access associated statistics (e.g., means, standard deviations, p-values, and fold changes calculated when comparing the biomarker in cases vs controls) in order to conduct or complement future meta-analyses. Finally, the API handles advanced database queries for extracting biomarker information that can be used to conduct complementary bioinformatics analyses as outlined by Zhang *et al*. in the context of colorectal cancer.

**Figure 3.**
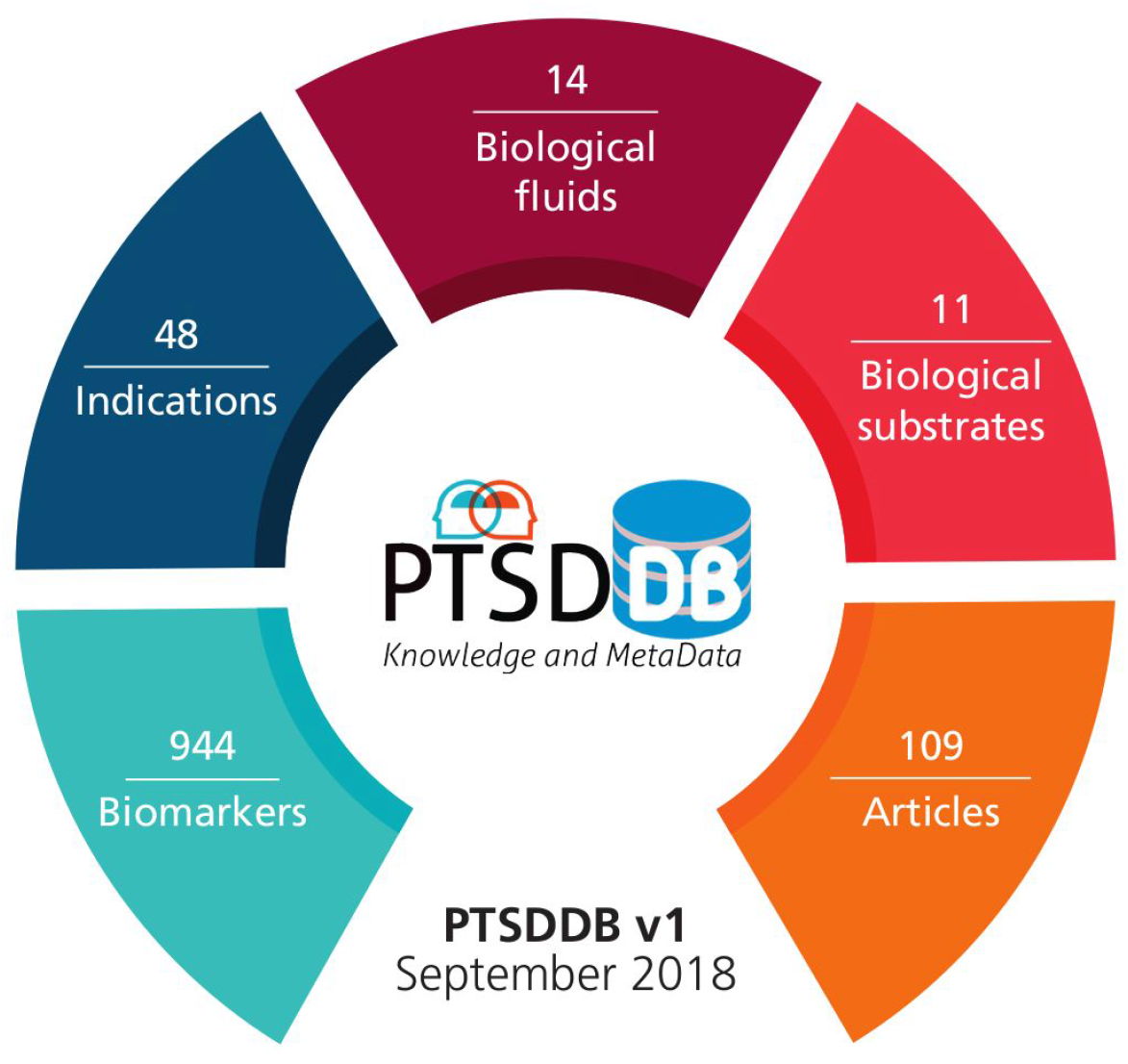
Content of the current version of PTSDDB: the database contains 109 articles, 944 biomarkers, 48 indications or distinct manifestations of PTSD, 14 biological fluids, and 11 substrates in which the biomarkers were tested.

## Conclusion

The PTSDDB organizes knowledge in the field of PTSD to provide a landscape view of the literature, bringing together results from different studies so that researchers can quickly and critically evaluate results of single biomarkers, understand how they were measured and in what population, and determine whether results are concordant to one another. This first version of the PTSDDB required a time-consuming effort that involved curating and harmonizing information from biomarker studies and related literature and storing this information in a database. In the future, this resource is planned to be integrated in Brain Commons (https://www.braincommons.org), a big data cloud-based platform for computational discovery designed with user-friendly tools for the research community. Per our knowledge, this paper describes the first resource designed to catalogue biomarker knowledge and metadata in PTSD. In summary, the PTSDDB is complemented with a comprehensive web application that provides interactive visualizations and tools to query the catalogued knowledge as well as advanced functionalities that can provide useful data for further statistical and bioinformatics analyses.

## Acknowledgements

We would like to thank the curators involved in the extraction of metadata from the articles included in the database.

