## Supplementary File for "PTSD Biomarker Database: Deep Dive Meta-database for PTSD Biomarkers, Visualizations, and Analysis Tools"

### Supplement

#### Articles included in the beta version of PTSDDB

*10200738 10202571 10715359 11847483 12114005 12207110 12562397 12798257 14744465 15026269 15199367 15289280 15685253 16325152 16585438 16647142 16891590 16934764 17853400 17884095 17888517 18037027 18055084 18277217 18295412 18342888 19107725 19269131 19393990 19394765 19409951 19576571 19628221 19751939 20097247 20298515 20439746 20488551 20621148 20643177 21078706 21145962 21196013 21208596 21350482 21508514 21508515 21536970 21714072 21906725 22137507 22867760 23319005 23379997 23650250 23701537 23774996 24035186 24362070 24485499 24576974 24661442 24704382 24709369 24759737 24929195 24952092 25080589 25106657 25129579 25311155 25345735 25706090 25745955 25754082 25800148 25827033 25848535 25867994 26074844 26112329 26120081 26120806 26151924 26160204 26174795 26277035 26305478 26324104 26361058 26447678 26481549 26632874 26649946 26673150 26714394 26751122 26894484 26923850 26997371 26999419 27025668 27038412 27193345 27479108 27716956 27886370 28521179 9137116*

**Supplement Text 1**. List of the 109 PubMed identifiers corresponding to articles included in the beta version of the database. The current list of articles can be accessed by querying the following API: https://ptsd.scai.fraunhofer.de/api/pubmed/all

### Survey of PTSDDB visualizations

1. **Biomarker overview** (https://ptsd.scai.fraunhofer.de/frequencies)

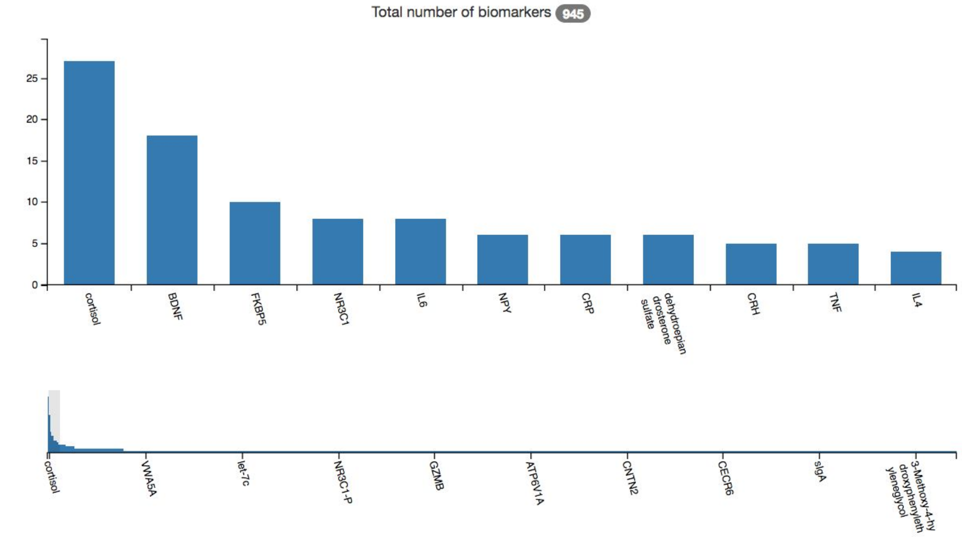


**Supplementary Figure 1.** Frequencies of the different biomarkers from the curated articles and included in PTSDDB.

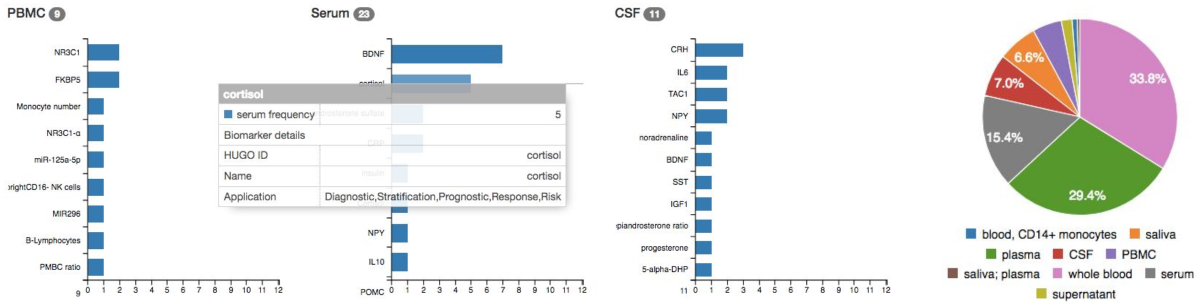


**Supplementary Figure 2.** Frequencies of the biomarkers by biofluids in which they were measured (left). These interactive visualizations offer more details of each biomarker when users hovers the bars. Biomarker frequencies by biological substrate and findings are represented as pie charts (right).

#### 2. Direction of change by biomarker

(https://ptsd.scai.fraunhofer.de/relative_changes)

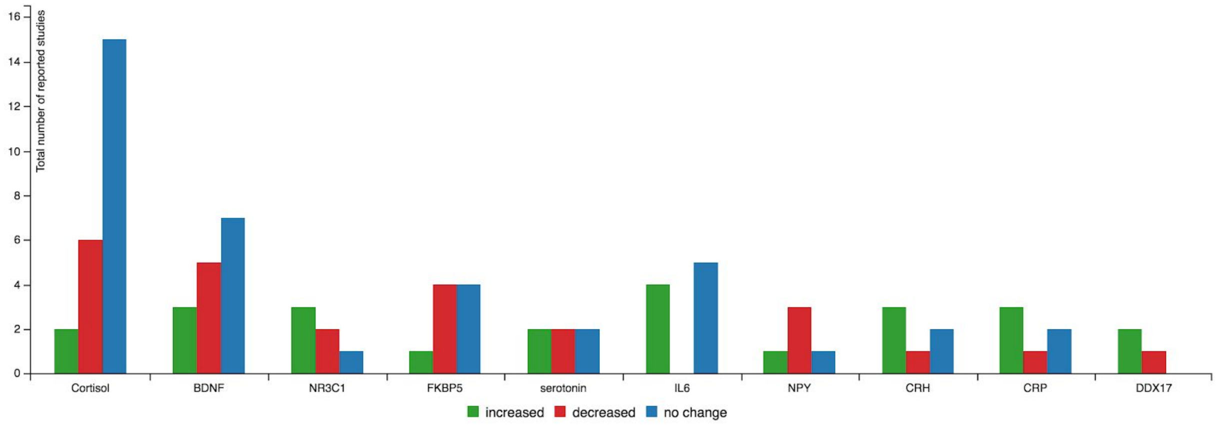


**Supplementary Figure 3.** Findings for the top ten studied biomarkers by valence of finding, representing the contradictory evidence found in the literature. Users can select other biomarkers and specifically visualize relative changes and the PubMed identifiers reporting those findings for a given biomarker.

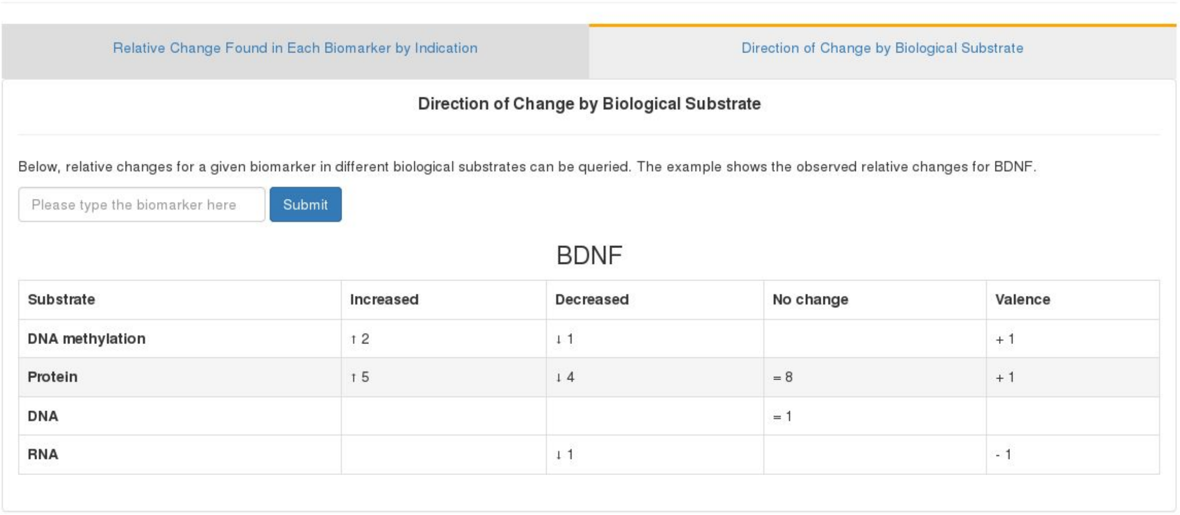
Relative Change Found in Each Biomarker by Indication

Direction of Change by Biological Substrate

Direction of Change by Biological Substrate

Below, relative changes for a given biomarker in different biological substrates can be queried. The example shows the observed relative changes for BDNF.

Please type the biomarker here

Submit

BDNF

| Substrate | Increased | Decreased | No change | Valence |
|---|---|---|---|---|
| DNA methylation | ↑ 2 | ↓ 1 | | + 1 |
| Protein | ↑ 5 | ↓ 4 | = 8 | + 1 |
| DNA | | | = 1 | |
| RNA | | ↓ 1 | | - 1 |

**Supplementary Figure 4.** Direction of change by biological substrate for a given biomarker. This interactive table presents the biomarker findings for different biological substrates (e.g., RNA, metabolite, protein, etc).

#### 3. Literature analysis (https://ptsd.scai.fraunhofer.de/literature)

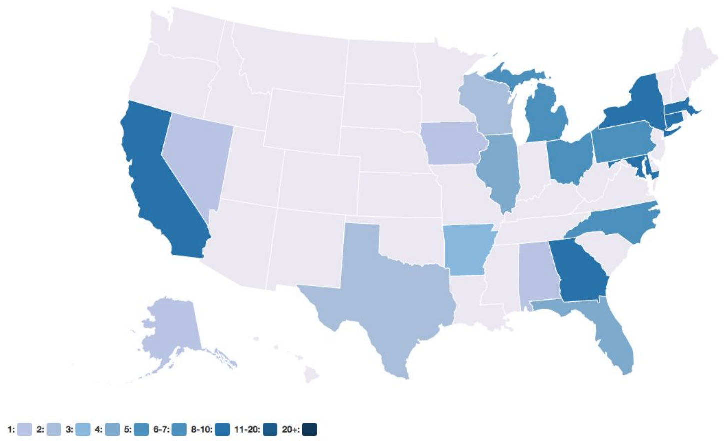


**Supplementary Figure 5.** Geographical distribution for the american authors in the articles included in PTSDDB. Additionally, a world map presents the full spectrum of countries involved in the articles curated (*Figure 2C* in the manuscript). This interactive view allows the identification of hub centers in the area of PTSD biomarker research.

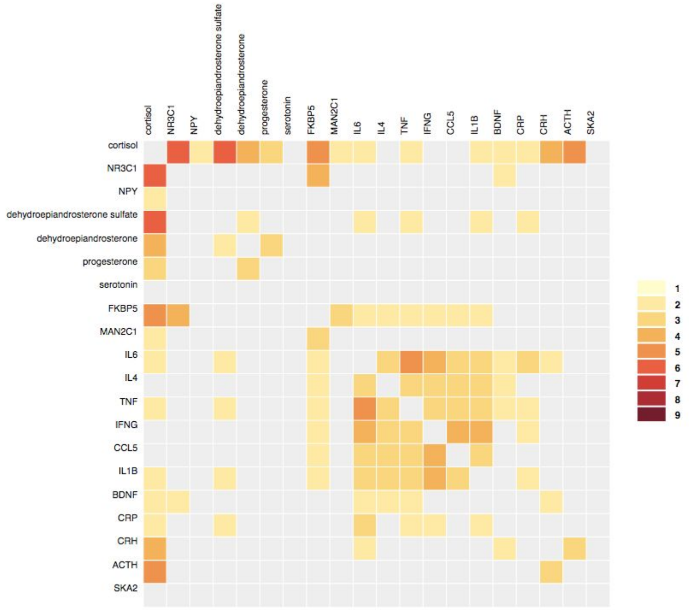


**Supplementary Figure 6.** Heatmap presenting the frequency in which pairwise-combination of biomarkers are studied together in the same publication. The heatmap can be interactively explored and allows for the clustering of biomarkers; in the example, cortisol emerges as the most commonly studied biomarker in combination with others. Moreover, we implemented two complementary heatmaps: 1) presenting the combination of biological substrates used in the extracted studies and 2) the combinations of biological substrates and bio-fluid utilized.

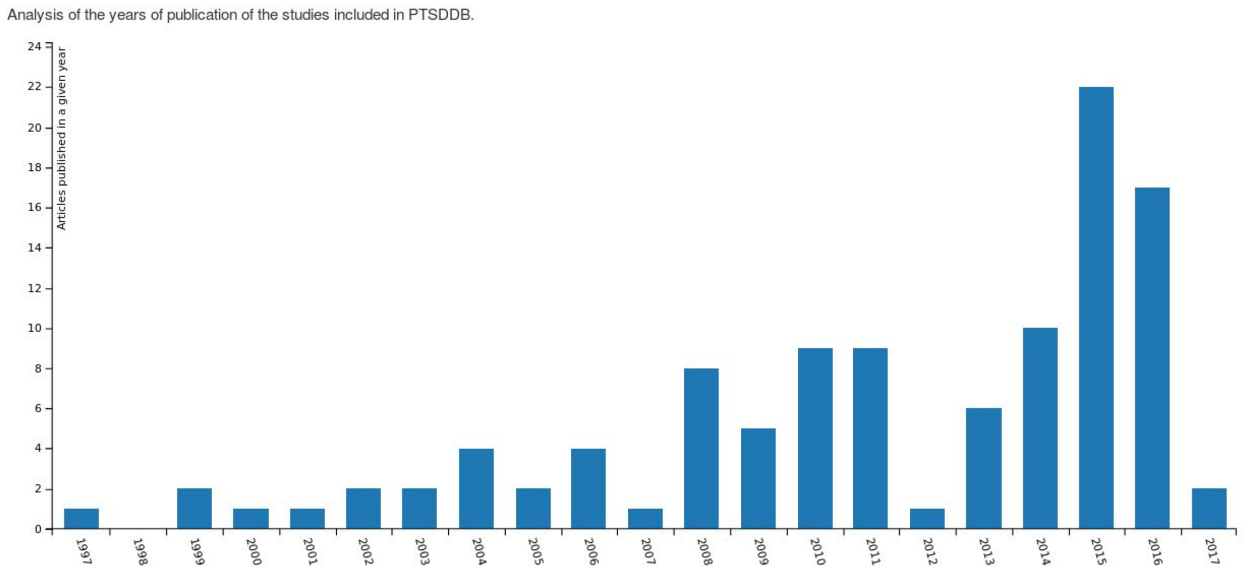


**Supplementary Figure 7.** Publication years of the articles included in PTSDDB.

#### 4. Study metadata (https://ptsd.scai.fraunhofer.de/study)

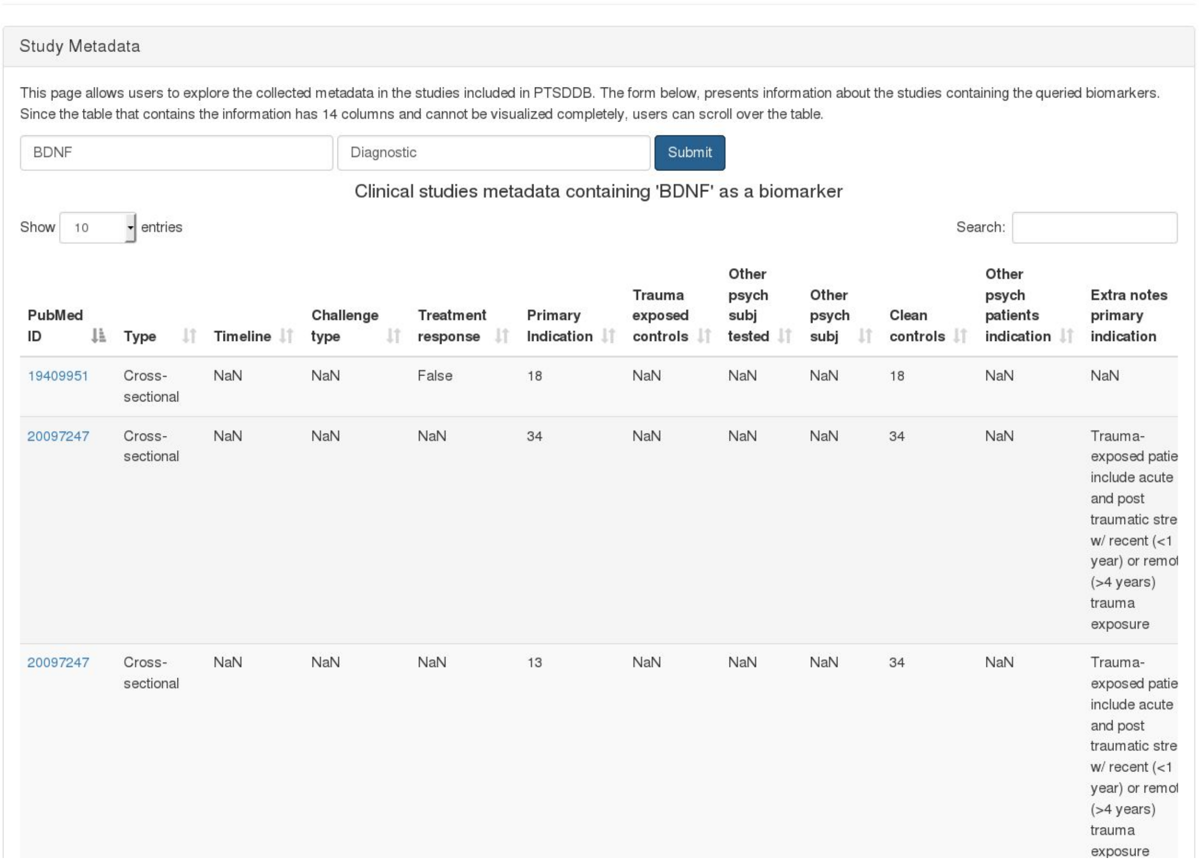

Study Metadata

This page allows users to explore the collected metadata in the studies included in PTSDDB. The form below, presents information about the studies containing the queried biomarkers. Since the table that contains the information has 14 columns and cannot be visualized completely, users can scroll over the table.

BDNF | Diagnostic | Submit

Clinical studies metadata containing 'BDNF' as a biomarker

Show 10 entries | Search:

| PubMed ID | Type | Timeline | Challenge type | Treatment response | Primary Indication | Trauma exposed controls | Other psych subj tested | Other psych subj | Clean controls | Other psych patients indication | Extra notes primary indication |
|---|---|---|---|---|---|---|---|---|---|---|---|
| 19409951 | Cross-sectional | NaN | NaN | False | 18 | NaN | NaN | NaN | 18 | NaN | NaN |
| 20097247 | Cross-sectional | NaN | NaN | NaN | 34 | NaN | NaN | NaN | 34 | NaN | Trauma-exposed patie include acute and post traumatic stre w/ recent (<1 year) or remot (>4 years) trauma exposure |
| 20097247 | Cross-sectional | NaN | NaN | NaN | 13 | NaN | NaN | NaN | 34 | NaN | Trauma-exposed patie include acute and post traumatic stre w/ recent (<1 year) or remot (>4 years) trauma exposure |

**Supplementary Figure 8.** The study metadata allows users to explore the collected metadata in the studies included in PTSDDB. The form above the page enables users to select biomarkers optionally filtered by their application to summarize the information about each clinical study that evaluated this biomarker.

#### 5. Database statistics and API (https://ptsd.scai.fraunhofer.de/overview)

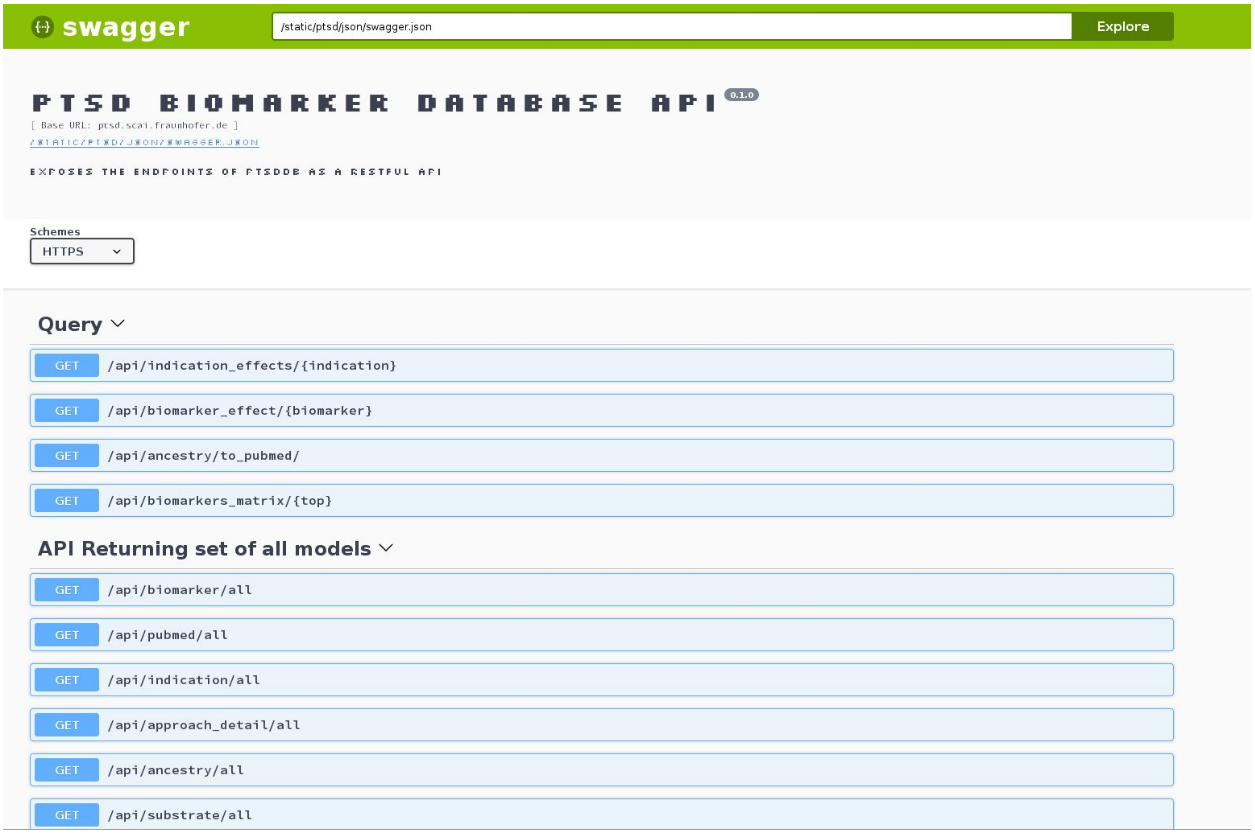


**Supplementary Figure 9**. The PTSDDB Swagger UI presents a user-friendly and interactive API visualization allowing developers and end consumers to interact with the API endpoints. This view is available at https://ptsd.scai.fraunhofer.de/swagger-ui.
